# Oral clade C SHIV challenge models to study pediatric HIV-1 infection by breastmilk transmission

**DOI:** 10.1101/545699

**Authors:** Koen K.A. Van Rompay, Alan D. Curtis, Michael Hudgens, Ryan Tuck, Neelima Choudhary, Maria Dennis, Ria Goswami, Ashley N. Nelson, Jeffrey Lifson, Ruth M. Ruprecht, George M. Shaw, Sallie R. Permar, Kristina De Paris

## Abstract

Nonhuman primate (NHP) models are invaluable for HIV pathogenesis, intervention and cure studies. To enhance the translational potential of NHP HIV vaccine studies, clinically relevant R5-tropic, tier 2 neutralization sensitive, and mucosally transmissible simian-human immunodeficiency viruses (SHIVs) have been designed. Towards our goal of developing vaccines to prevent breastmilk transmission of HIV, we evaluated virological outcomes of three distinct SHIVs in repeated weekly oral exposure regimens in infant rhesus macaques. The selected strains SHIV-1157ipd3N4, SHIV-1157(QNE)Y173H, and SHIV CH505 375H.dCT express a clade C HIV Env, the clade most prevalent in regions with high pediatric HIV infections.

All three SHIVs were orally transmissible. However, compared to the pediatric SIV_mac251_ model, SHIV replication was more attenuated. Some animals exposed to weekly low-dose (20 TCID_50_) SHIV-1157ipd3N4 had transient viremia blips associated with lack or delayed seroconversion. Animals with acute viremia ≥10,000 viral RNA copies/ml seroconverted. This finding was reminiscent of an earlier study suggesting to continue challenges until a threshold of ≥10,000 viral RNA copies/ml is surpassed to achieve seroconversion, one criterion of HIV diagnosis. All animals exposed weekly with higher doses (>10^4^ TCID_50_) of SHIV-1157(QNE)Y173H or SHIV CH505 375h.dCT developed persistent infection with high peak viremia and seroconverted. Chronic viremia varied widely in all three SHIV infection models. Thus, although R5 clade C SHIVs are excellent tools to study prevention of virus acquisition, secondary virological outcomes of vaccine efficacy (e.g. risk of infection per exposure, modulation of peak viremia, viral set point) will require careful consideration and validation in SHIV challenge models.

**IMPORTANCE:** The development of an effective HIV vaccine remains a top priority towards the goal of reducing the number of new HIV infections. Studies in nonhuman primate models of HIV infection have been instrumental in the preclinical evaluation of candidate vaccines. To assess the role of HIV Env-specific antibodies with broadly neutralizing or Fc-mediated effector function in these NHP models, the development and optimization of novel SHIV challenge models is required. We evaluate three different clade C HIV Env SHIVs for infectivity of infant macaques by the oral route. Our results demonstrate that SHIV-1157ipd3N4, SHIV-1157(QNE)Y173H, and SHIV CH505 375H.dCT can be orally transmitted and establish persistent infection in infant rhesus macaques, but chronic viremia levels vary widely. Therefore, SHIV infection models are most pertinent when prevention of systemic infection is the primary readout of vaccine efficacy, but secondary outcome measures of vaccine efficacy will require stringent criteria and extensive validation.

## INTRODUCTION

The successful implementation of antiretroviral therapy (ART) for HIV-infected women has resulted in a dramatic reduction of mother-to-child transmission of HIV-1 in the last two decades. Yet, globally, between 400 to 500 infants acquire HIV every day (1). The majority of these infections occur during the breastfeeding period. Limited access to ART in rural communities, HIV diagnosis late in pregnancy, gaps in linking antenatal care with postnatal mother and infant care, acute infection of the mother during the breastfeeding period, and lack of ART adherence continue to impede the prevention of HIV transmission via breast milk (2–9). Transmission of HIV can occur throughout the entire breastfeeding period, with a cumulative risk increase with every month of breastfeeding (10–13). In many resource-limited countries, breast milk remains a necessary choice of nutrition and passive immunity to protect the infant against the many other pathogens that are highly prevalent in these areas (6, 7, 14). Indeed, early weaning is associated with increased infant mortality (15–17), and the WHO recommends exclusive breastfeeding for 6-12 months for HIV-exposed infants (18). Therefore, the development of an HIV vaccine that can protect infants remains a high priority.

Importantly, whereas infants born to mothers with known HIV-positive status are tested at birth and immediately started on ART if HIV-positive, infants that acquire HIV by breastfeeding often go undiagnosed until routine HIV testing after the cessation of breastfeeding or when they develop clinical symptoms. Considering that the breastfeeding period coincides with the time of rapid development of both the immune system and the central nervous system, prolonged HIV replication prior to diagnosis may severely interfere with multiple aspects of normal development and impede immune reconstitution after ART initiation. Thus, to prevent long-term health sequelae, we need a deeper understanding of the factors controlling HIV pathogenesis and persistence in infants.

Animal models of pediatric HIV infection are an invaluable resource for addressing these questions. The oral SIV_mac251_ infection model in infant rhesus macaques simulates the most important aspects of HIV acquisition by breastfeeding in human infants (19). The pediatric SIV_mac251_ infection model has been extensively utilized to study early host responses (20–24), test new treatment strategies (25–27), and to optimize pediatric HIV vaccine strategies (28–32). To further increase the translational value of the pediatric oral SIV_mac251_ infection model, the single high-dose challenge has been replaced by a repeated weekly oral SIV_mac251_ exposure model to more closely resemble actual HIV exposures in human infants (33–35).

In HIV vaccine design, the induction of protective HIV Env-specific antibody responses has taken center stage. As passive immunization studies in NHP models have shown that broadly neutralizing antibodies (bnAbs) can prevent infection, one major goal is the development of vaccines that induce such bnAbs and could protect against the various distinct HIV clades circulating worldwide. In the RV144 HIV vaccine trial in human adults, however, antibodies with Fc-mediated effector function contributed to the protection against HIV acquisition (36). To evaluate HIV Env-specific antibody responses in nonhuman primate models, chimeric SIV/HIV Env (SHIV) challenge viruses and infection models are needed. Although multiple SHIVs carrying HIV-1 envelopes have been developed previously, these earlier SHIVs were not characteristic of clinically relevant transmitted/ founder (T/F) viruses and sometimes exhibited CXCR4 tropism (37, 38). The current study took advantage of ongoing pediatric vaccine studies in our groups that utilized different SHIVs as challenge virus. Towards the goal of developing a clinically relevant clade C SHIV infection model to study HIV pathogenesis as well as vaccine and cure strategies directed towards the pediatric population, we specifically tested SHIV-1157ipd3N4, SHIV-1157(QNE)Y173H, and SHIV CH505 375H.dCT. These SHIVs were selected because they were designed with an HIV env gene derived from a T/F virus of clade C, the prominent clade in sub-Saharan Africa where most pediatric HIV infections occur. Enhancing their clinical relevance, these SHIVs are CCR5-tropic, mucosally transmissible, and have tier 2 neutralization susceptibility (39–41). Building on our experience with the pediatric oral SIV_mac251_ infection model, we evaluated these SHIVs for their virologic profile in infant macaques infected by the oral route applying a repeated, weekly oral exposure regimen, and discuss the potential value of the different challenge models for vaccine efficacy studies in NHP models to better inform future clinical HIV vaccine trials.

## RESULTS

### Review of the infant macaque model of oral SIV_mac251_ infection

Nonhuman primate models of pediatric HIV infection have been developed and refined over the past 25 years (26, 42–44). The initial neonatal high-dose oral SIV infection model (20) was subsequently refined to simulate the repeated HIV exposure of human infants by breast milk by bottle-feeding SIV_mac251_ to neonatal macaques 3 times per day for 5 consecutive days (31). This regimen resulted in systemic SIV infection in >80% of infants within 2 weeks after the first exposure, but had the drawback that, as inoculations were all provided in a narrow time window, it was not possible to identify which inoculation induced infection (21, 28). We subsequently developed a repeated weekly oral SIV_mac251_ exposure model (33–35). The virological outcome of this repeated weekly challenge strategy was highly consistent; in studies spanning 3 years, the estimated risk of infection per exposure ranged from 0.23-0.25 (33–35). Animals exhibited high peak viremia (median: 4.8×10^7^ copies/ ml) and viremia persisted at high levels (at week 8 post-infection [PI]: median: 1.1×10^7^ copies/ ml) (Figure 1). This oral SIV_mac251_ infection model in infant macaques best recapitulates the outcome of HIV-infected human infants that progress rapidly to AIDS in the absence of ART.

**Figure 1:**
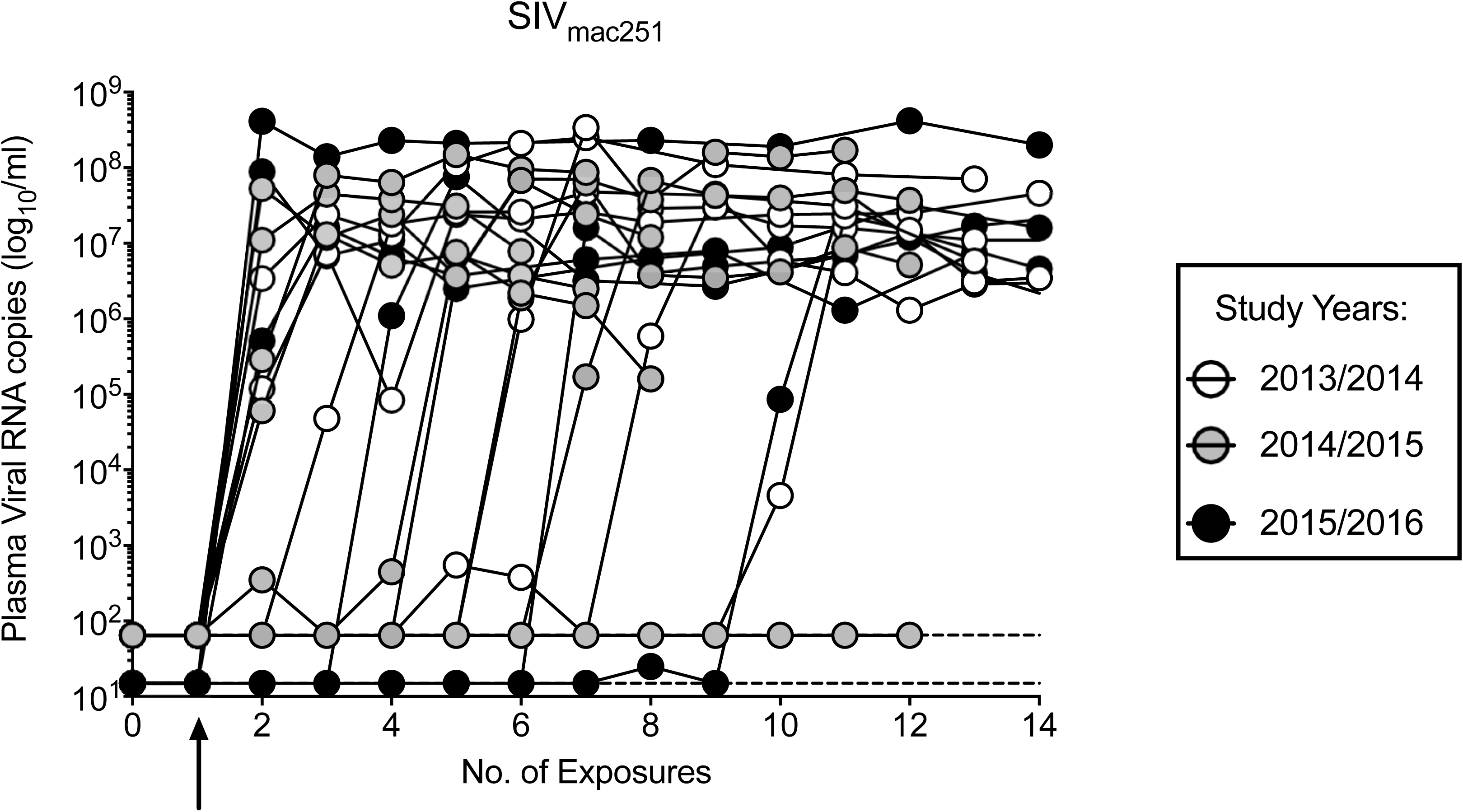
Plasma viremia of orally SIV_mac251_-infected infant macaques. Each curve represents a single infant macaque orally infected with SIV_mac251_ by a repeated weekly exposure regimen (n=21). The data represent a summary of three independent studies in three subsequent years, indicated by different shades as specified in the graph legend (33–35). Plasma viremia is graphed in relation to the number of viral exposures. Th earrow below the x-axis indicates the first viral exposure. Note that the lower detection limit (LOD) of the viral RNA PCR for the studies performed in 2013-2015 was 65 copies/ ml, but 15 copies/ml for the 2015/2016 study, and is indicated by thin dashed lines.

The reproducibility of the oral pediatric SIV_mac251_ infection model constitutes a fundamental value for the study of pediatric HIV. However, as with any animal model, and dependent on the purpose of the study, certain limitations may apply. In transmission studies, the evaluation of T/F viruses is hampered by the limited genetic diversity. Although SIV_mac251_ is an uncloned viral swarm, the genetic diversity within each stock is often low and variable among different SIV_mac251_ challenge stocks because of prior histories of propagation (22, 34, 45). In addition, due to significant differences in envelope between SIVmac (which is similar to HIV-2) and HIV-1, the use of the oral pediatric SIV_mac251_ infection model also has limited translational potential for HIV vaccine studies assessing the role of HIV Env-specific antibody responses in vaccine-mediated protection against HIV acquisition. Therefore, we aimed to develop novel oral SHIV C infection models. Analogous to the pediatric SIV_mac251_ infection model, we tested 3 different SHIVs in a repeated, weekly oral SHIV exposure model with the goal to achieve an infection risk of 25% to 50% per exposure in infant macaques. Virus was applied atraumatically in 1 ml volume to the oral mucosa and animals were considered systemically infected when plasma viral RNA was detected at two consecutive time points.

### A pediatric SHIV-1157ipd3N4 infection model

SHIV-1157i was one of the first SHIVs developed that contains a clade C HIV Env (46). We considered this SHIV as being potentially of particular clinical relevance, because HIV-1157 was isolated from an HIV-infected African infant. The original SHIV-1157i underwent several adaptations via *in vivo* serial passage in macaques to increase pathogenicity (41, 46, 47). Furthermore, the SIV_mac239_ LTR was engineered to contain additional NF-κB sites (48). We selected the clone SHIV-1157ipd3N4 for our studies because it was able to infect adult macaques by multiple mucosal routes, including the oral route (49).

Based on data in adult macaques (49), we tested a dose of 800 TCID_50_ SHIV-1157ipd3N4 in infant macaques. All infant macaques (n=3) became infected after a single oral exposure (Figure 2A). We then exposed a second group of 4 infant macaques to only 10 TCID_50_ of SHIV-1157ipd3N4. Plasma viremia became detectable in 2 of the 4 infants after 3 challenges and these 2 infants received no further oral SHIV exposures (Figure 2B). The remaining 2 infant macaques developed viremia after the dose for the 4^th^ exposure was increased to 50 TCID_50_ of SHIV-1157ipd3N4 (Figure 2B).

**Figure 2:**
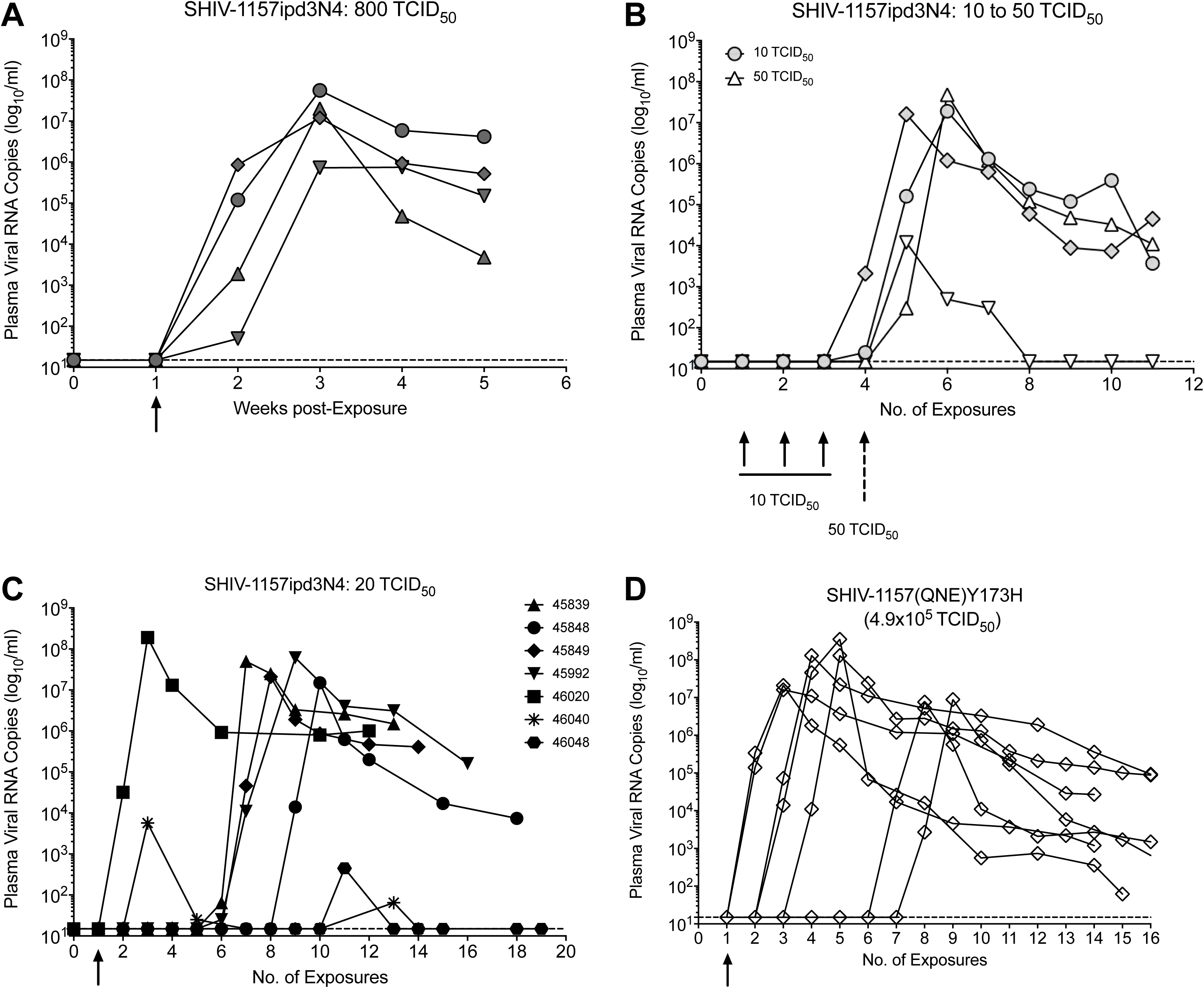
SHIV-1157ipd3N4 virus titration and viremia. Infant macaques were orally exposed to various doses of SHIV-1157ipd3N4. *Panel A:* Four infant macaques orally challenged with 800 TCID_50_ of SHIV-1157ipd3N4 became infected after a single exposure. *Panel B:* Exposure of infant macaques with 10 TCID_50_ of SHIV-1157ipd3N4 resulted in systemic infection after 3 oral exposures in 2 of 4 animals, with the remaining 2 animals becoming infected after a single, subsequent exposure to 50 TCID_50_ of SHIV-1157ipd3N4. *Panel C:* Plasma viremia in 7 infant macaques exposed once weekly to 20 TCID_50_ of SHIV-1157ipd3N4. Animals with transient viremia are indicated by grey symbols and dashed lines. *Panel D:* Plasma viremia in infant macaques (n=7) infected with the related virus SHIV-1157(QNE)Y173H applying the same repeated oral exposure regimen as in Panel C. Each curve in each of the panels represents an individual animal. The LOD (15 copies/ ml) for the viral RNA PCR is indicated by thin dashed lines. Arrows below the x-axis in Panels A, C, and D indicate the first exposure.

Based on this initial virus titration, we proceeded with a weekly oral exposure regimen of 20 TCID_50_ in 7 infant macaques starting at week 6 of age (50). Five of 7 SHIV-1157ipd3N4 infected infant macaques developed high peak viremia (>10^6^ viral RNA copies/ ml) that subsequently declined at variable levels (Figure 2C). The risk of infection per oral exposure (0.21) in these 5 infant macaques was comparable to the oral SIV_mac251_ infection model (50). The other two infants (#46040, #46048) had only transient viremia. One of these infant macaques tested positive for 5700 viral RNA copies /ml plasma after the first exposure and viral RNA was still detectable at the next time point, although at very low levels of only 25 copies/ ml (Figure 2C). Therefore, based on our criteria that an animal would be considered infected when viral RNA was detectable at two consecutive time points, this animal received no further challenges. In contrast to these 6 infants, the remaining animal (#46048) remained viral RNA negative after 7 exposures to 20 TCID_50_ of SHIV-1157ipd3N4 (Figure 2C). Therefore, the dose was increased to 40 TCID_50_ of SHIV-1157ipd3N4, and a single plasma viral RNA “blip” of 450 copies /ml was detected after 2 challenges (Figure 2C); viral RNA was not detected at any later time point. Consistent with our infection criteria, we considered this animal uninfected after a total of 13 (7×20 TCID_50_ plus 6×40 TCID_50_) weekly SHIV-1157ipd3N4 challenges (50). Reduced viremia or lack of persistent viremia, as previously reported, was not associated with protective MHC or TRIM5 genotypes (50), but instead was likely the result of the very low exposure dose, a dose 100x lower than the dose in the repeated oral SIV_mac251_ challenge model. This conclusion is indirectly supported by the challenge outcome of infant macaques that received weekly oral challenges of 4.9×10^5^ TCID_50_/ml SHIV-1157(QNE)Y173H, a derivative of SHIV-1157 with 3 mutations in the V2 region. All animals developed high peak viremia (Figure 2D). Chronic viremia of SHIV-1157(QNE)Y173H-infected infant macaques, similar to the SHIV-1157ipd3N4 animals with high peak viremia, however, was also highly variable (Figure 2D).

The lack of persistent viremia in 2 of the 7 SHIV-1157ipd3N4-exposed infants was consistent with a lack of seroconversion in animal #46048, and delayed development of plasma gp120-specific antibody responses in the other animal (Figure 3). In contrast, all infants with persistently high viremia in the acute phase seroconverted within 2 to 4 weeks of peak viremia (Figure 3). These data implied that seroconversion, a parameter clinically used to confirm HIV infection in humans, requires a certain threshold of viremia.

**Figure 3:**
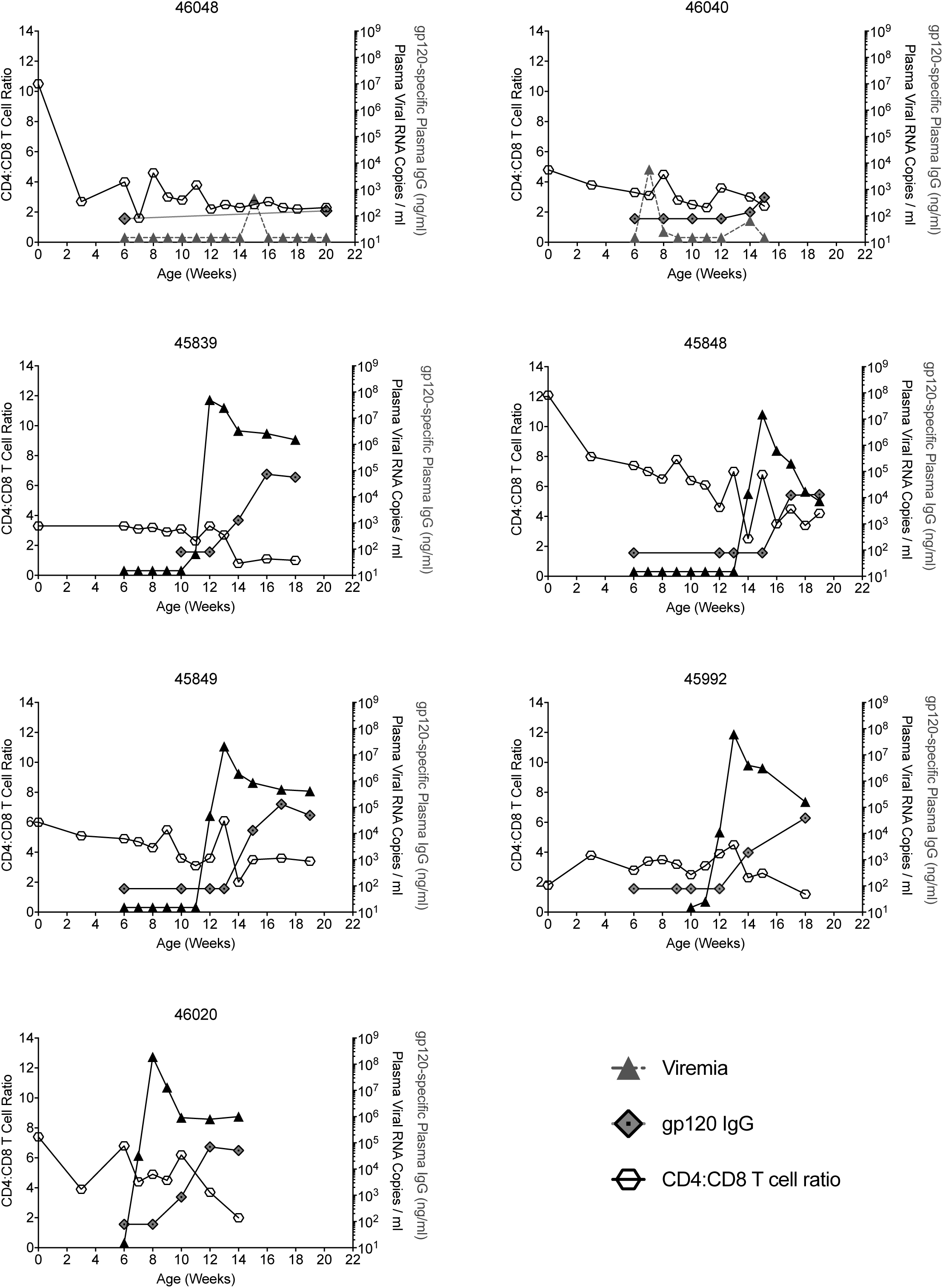
Correlation between plasma viremia, development of plasma gp120-specific IgG antibodies and the ratio of CD4:CD8 T cells. Each graph represents an individual animal orally infected with 20 TCID_50_ of SHIV-1157ipd3N4 (see Figure 2C). The x-axis refers to the age of the animal in weeks. Virus exposure began at 6 weeks of age and plasma viremia is illustrated by the black line with black triangles. The induction of gp120-specific plasma IgG antibodies in response to infection is indicated by the line with grey diamond symbols, with antibody measurements starting at week 6, the time of the first oral SHIV exposure. To document the normal developmental decline in the CD4:CD8 T cell ratio (open hexagons) independent of viral infection, the CD4:CD8 ratio is displayed from birth (week 0) to the study end.

In addition to seroconversion, we also assessed CD4^+^ T cell loss. However, due to the normal developmental decline of CD4^+^ T cells and the concurrent increase in CD8^+^ T cells in both human and rhesus infants after birth, CD4^+^ T cell loss in peripheral blood is not a reliable clinical marker of early SIV or SHIV infection (28, 51-53). Indeed, the peripheral blood CD4:CD8 T cell ratio continuously declined in most animals prior to SHIV-1157ipd3N4 infection (Figure 3). Compared to week 0 of infection, defined as the last viral RNA PCR-negative time point for each animal, the CD4:CD8 T cell ratio exhibited a further median decline of 35% (range 0-50%) or 40% (range 0-70%) by week 3 or 8 PI, respectively (Figure 3).

### An oral SHIV CH505 375H.dCT infection model in infant macaques

The HIV Env in SHIV CH505 375H.dCT, similar to SHIV-1157ipd3N4, is also derived from a clade C, R5-tropic T/F virus with a tier 2 neutralization phenotype, although it was isolated from an adult HIV-infected patient who developed broadly neutralizing activity (54). Recognizing the differences in the interactions of HIV Env or SIV Env with the rhesus CD4 molecule, the HIV *env* gene was mutated at position 375 to enhance binding to rhesus CD4, thereby increasing pathogenicity in rhesus macaques (40). In adult macaques, CH505 375H.dCT was infectious by the intravenous (IV), intravaginal (IVAG), and intra-rectal (IR) route (Dr. Shaw, pers. comm.).

To determine an oral infectious CH505 375H.dCT dose in infant macaques, we titrated the virus *in vivo* starting with a 1:80 dilution (equal to 8.5×10^4^ TCID_50_/ml) of the virus stock. The first animal remained uninfected after 6 weekly oral exposures to CH505 375H.dCT (Figure 4A). The oral challenge dose was then increased weekly (1:40, 1:20, 1:10, 1:5, and 1:2) until the animal eventually became infected after 5 exposures to undiluted virus (6.8×10^6^ TCID_50_/ml; Figure 4A). A second animal, however, became infected after a single exposure to 8.5×10^4^ TCID_50_/ml of CH505 375H.dCT, whereas two additional animals remained uninfected after two challenges, but became viremic after a single exposure to 17×10^4^ TCID_50_/ml (Figure 4B).

**Figure 4:**
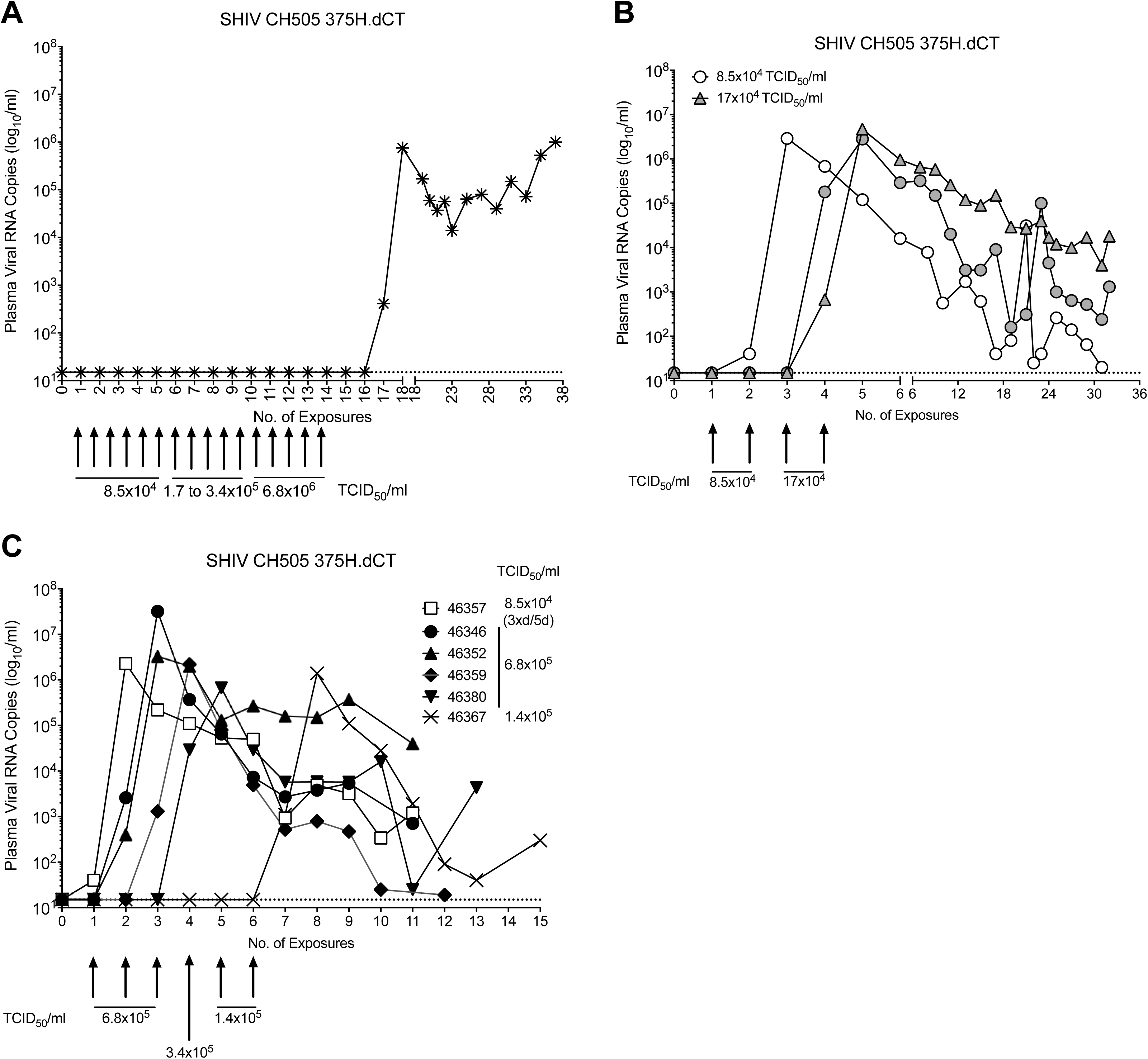
In vivo infectivity of SHIV CH505 375H.dCT in infant macaques. *Panel A*: Oral SHIV CH505 375H.dCT dose escalation in a single animal to establish systemic infection. *Panel B*: Plasma viremia of 3 infant macaques exposed once weekly to 8.5×10^4^ or 17×10^4^ TCID_50_/ml of SHIV CH505 375H.dCT at the times indicated by arrows underneath the x-axis. *Panel C*: Plasma viremia of 6 infant macaques that were exposed to SHIV CH505 375H.dCT 3x/day for 5 days (each dose at 8.5×10^4^ TCID_50_/ml), and subsequently underwent a once weekly oral challenge with SHIV CH505 starting at 6.8×10^5^ TCID_50_/ml dilution. In each panel, the x-axis displays the number of oral virus exposures with black arrows indicating the time and dose. The LOD (15 copies/ ml) of viral RNA detection is indicated by thin dashed lines.

A second group of infant macaques (n=6) at 4 weeks of age was exposed via bottle-feeding 3 times daily at a dose of 8.5×10^4^ TCID_50_/ml for 5 consecutive days, a regimen we had successfully used with SIV_mac251_ to simulate repeated exposure to HIV by breastfeeding (21, 28). However, only one of the infants became infected using this exposure regimen and had a low plasma viremia of 40 copies/ml at week 1 after the first exposure (Figure 4C). Therefore, and based on the challenge outcome in the first group of infant macaques, we decided to challenge the uninfected animals of the second group once weekly with 6.8×10^5^ TCID_50_/ml SHIV CH505 375H.dCT. Applying this regimen, 2 animals became infected after 1 exposure, and 1 animal each at 2 or 3 exposures (Figure 4C). We then increased the exposure dose and the last animal became infected after two oral exposures of 3.4×10^6^ TCID_50_/ml SHIV CH505 375H.dCT (Figure 4C). Independent of the dose required to infect the animals, all animals developed high peak viremia, an outcome similar to the oral SIV_mac251_ infection in infant macaques. In contrast to SIV_mac251_ infection, but similar to the infections with SHIV-1157ipd3N4 or SHIV-1157(QNE)Y173H, viremia became highly variable during the chronic phase without reaching a specific set point (Figure 4). Although the follow-up time was short (<10 wks PI), viremia appeared to fluctuate more compared to the SHIV1157ipd3N4-infected animals (see Figure 2). To confirm persistent viral infection and the establishment of a viral reservoir, we analyzed lymph nodes for virally infected CD4^+^ T cells. Consistent with SIV and HIV pathogenesis studies, virally-infected cells were predominantly found in B cell follicles of SHIV CH505 375H.dCT and SHIV-1157ipd3N4-infected macaques (Figure 5).

**Figure 5:**
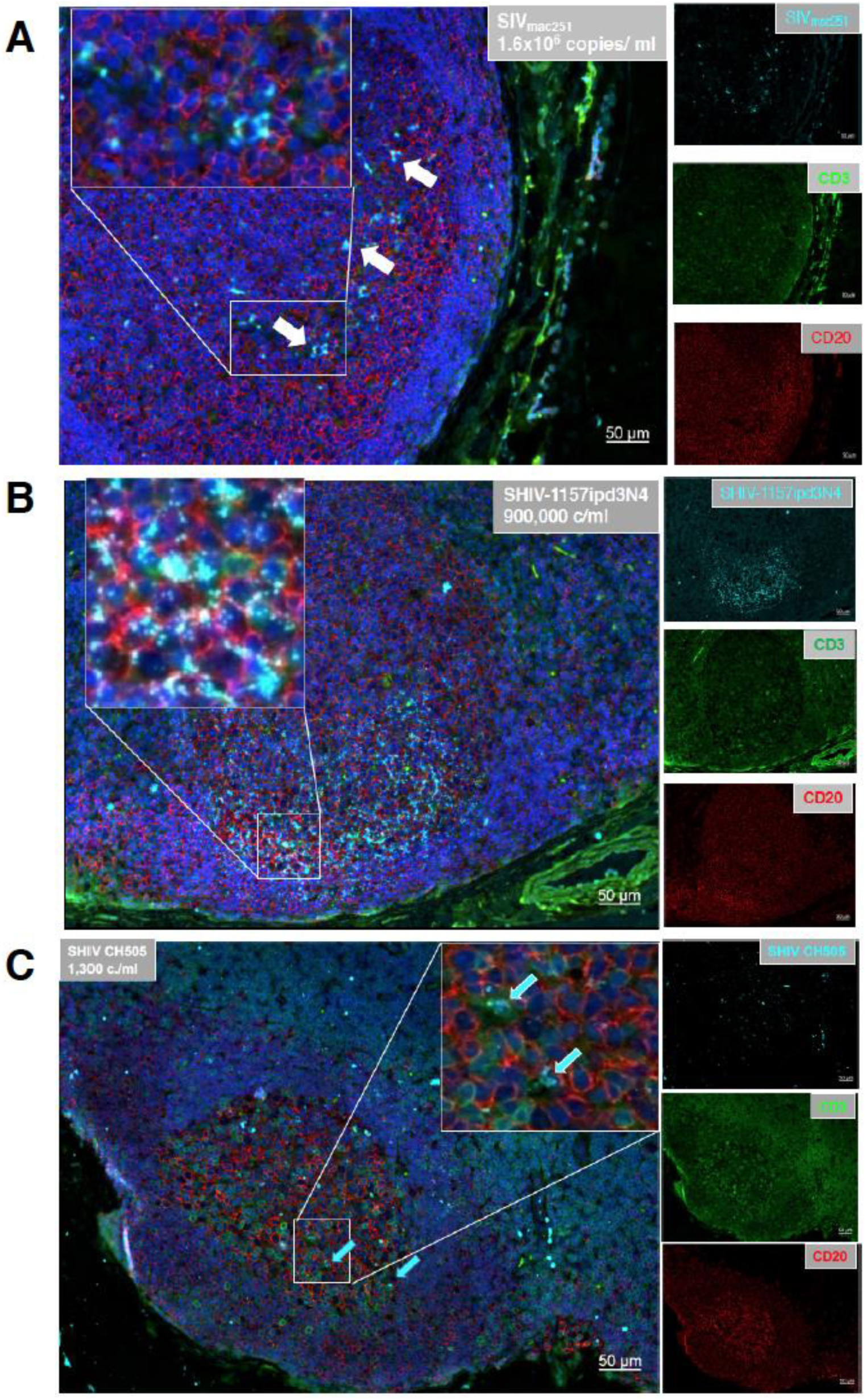
Virus detection in lymph nodes of virally-infected infant macaques. Panels A to C show representative examples of lymph node sections from SIV_mac251_ (A), SHIV-1157ipd3N4 (B), or SHIV CH505 375H.dCT (C)-infected infant macaques, respectively. Sections were stained with the nuclear marker DAPI (dark blue) to identify cells, and with antibodies specific for CD3 (green) and CD20 (red). Virally-infected cells were identified by in-situ hybridization (cyan). To better visualize the virally-infected cells, we magnified a specific region (white box) in each image. Each panel consists of a larger image with the overlay of all markers and 3 smaller side panels of the same field for each individual channel. Arrow colors correspond to the indicated marker. The large image has a scale bar in the lower right corner.

As observed for SHIV-1157ipd3N4-infected infant macaques with high acute viremia, seroconversion occurred within the first 2-4 weeks of SHIV CH505 375H.dCT infection (Figure 6). The magnitude of plasma gp120-specific IgG antibodies appeared to be higher in SHIV CH505 375H.dCT (range from 10^4^-10^6^ ng/ml) compared to SHIV-1157ipd3N4 (maximum of 10^5^ μg/ml) infected infant macaques, but as we only followed the infection for 8 weeks PI, it is possible that maximum plasma gp120-specific IgG antibody levels in some animals had not been reached at this time point.

**Figure 6:**
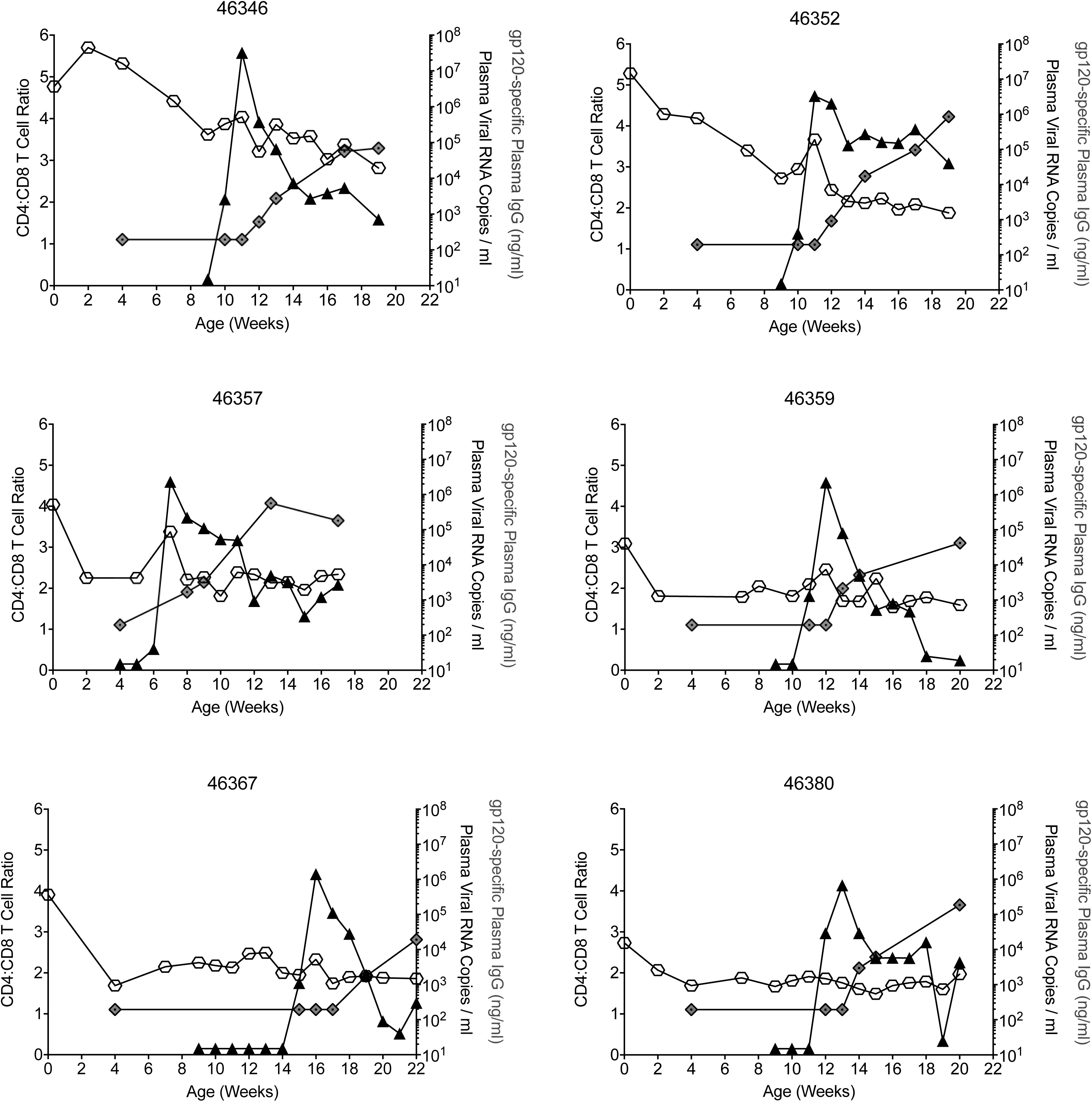
SHIV CH505 375H.dCT plasma viremia in relation to plasma gp120-specific IgG antibodies and the CD4:CD8 T cell ratio. The figure is designed analogous to Figure 3, except that the oral challenge with CH505 375H.dCT was started at week 9 of age (see Figure 4C).

**Figure 7:**
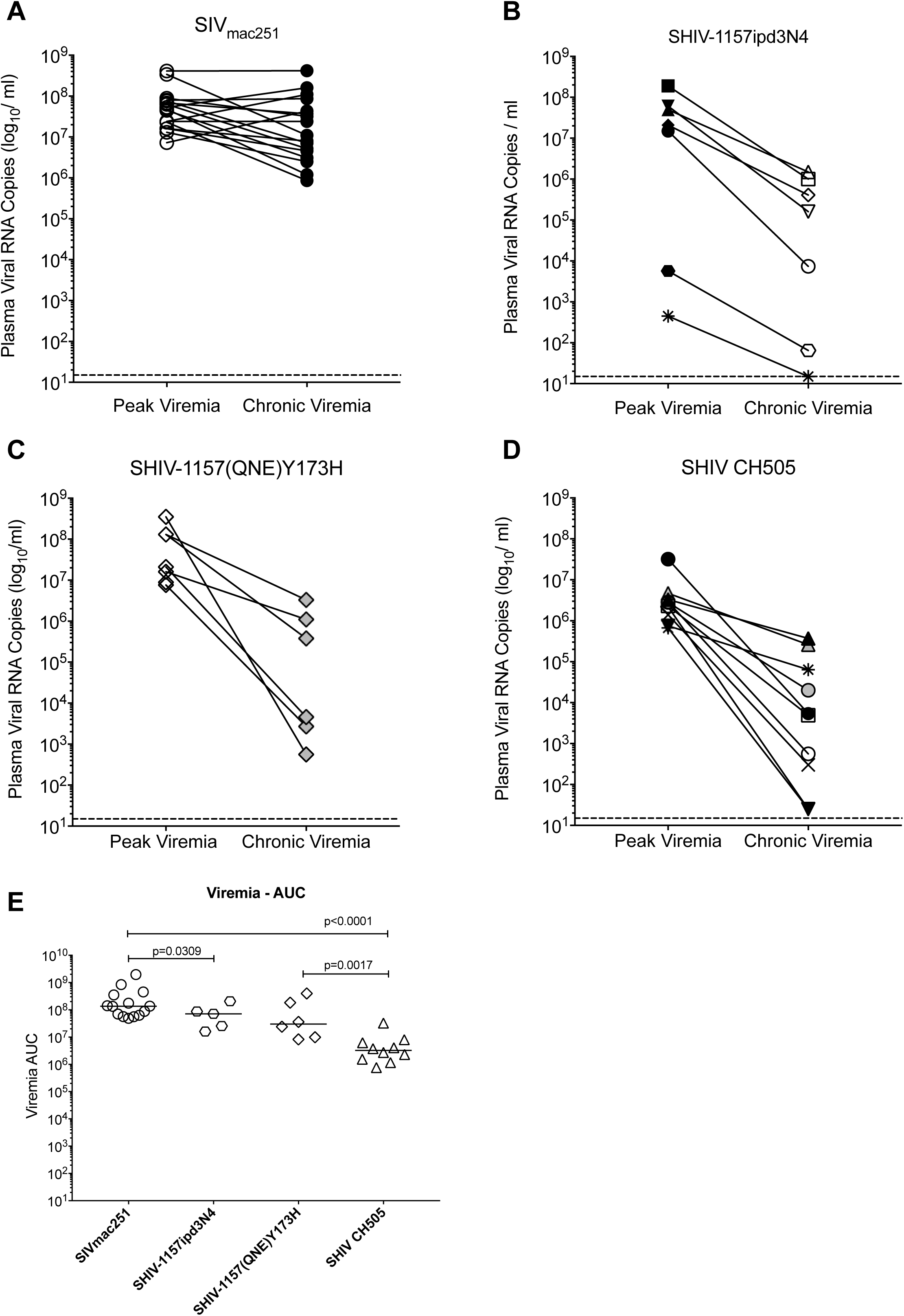
Peak and chronic viremia in infant macaques infected with SIV_mac251_ or various SHIVs. *Panels A-D* display the peak viremia and early chronic phase viremia (week 8 PI) for SIV_mac251_ (A), SHIV-1157ipd3N4 (B), SHIV-1157(QNE)Y173H (C), and SHIV CH505 375H.dCT (D)-infected infant macaques. Each line represents an individual animal. The animals in Panels B and D have the same symbols that we reused in the corresponding Figures 2C and 4C, respectively. Data in Panel A and Panel C refer to the animals in Figure 1 and Figure 3D, respectively. The thin dashed line corresponds to the LOD of the viral RNA PCR. *Panel E* shows the comparative AUC (weeks 0 to 8 PI) plasma viremia data for the four different SHIVs tested.

We also observed an age-related decrease in the CD4:CD8 T cell ratio prior to oral SHIV CH505 375H.dCT exposure (Figure 6). While the CD4:CD8 T cell ratio continued to decline in 2 of the 6 SHIV CH505-infected animals, the CD4:CD8 T cell ratio remained relatively stable in the remaining 4 infants and by 8 weeks PI, the median CD4:CD8 T cell ratio had decreased by 11% (range: 0% to 23%; Figure 5). There was no apparent correlation between the magnitude of virus replication and the decline of the CD4:CD8 T cell ratio in either of the SHIV infection models. However, due to the short follow-up time, we could not assess whether continuous virus replication would have resulted in a more pronounced CD4^+^ T cell decline over time.

## DISCUSSION

The goal of the current study was to determine the potential of different SHIVs as challenge viruses in pediatric NHP vaccine efficacy studies. The different SHIVs, SHIV-1157ipd3N4, SHIV-1157(QNE)Y173H, and SHIV CH505 had been selected based on their potential clinical relevance and translatability of the results to human clinical studies.

In previous studies evaluating pediatric vaccine candidates intended to prevent breast milk transmission of HIV in human infants, we employed the repeated oral SIV_mac251_ exposure model in infant macaques (33–35). Consistent behavior of the SIV_mac251_ challenge virus is a strength of that model; in multiple studies the risk of infection per SIV_mac251_ exposure, peak viremia, as well as chronic viremia were highly similar among all animals across all studies (33–35). These findings provided the basis for the development of the repeated oral SHIV exposure models. Our results demonstrate that all three SHIVs could infect infant macaques via repeated oral exposure. Therefore, all three SHIVs qualify as challenge models to evaluate pediatric vaccines for their ability to protect against virus acquisition, the ultimate goal of an HIV vaccine.

NHP models for HIV/AIDS should reflect the biology of HIV-1 transmission in humans as closely as possible to maximize the predictive value of NHP vaccine safety and efficacy for testing of HIV vaccine candidates in humans. People enrolled in such trials may not have the same close follow-up that can be applied in NHP studies, where viral RNA loads are measured once or twice a week. As a consequence, vaccine efficacy in clinical trials is assessed mostly by seroconversion. In HIV-exposed infants, the presence of maternal antibodies may prevent the use of seroconversion as a clinical read-out. The generation of specific antiviral antibodies takes several weeks in HIV-infected humans as well as in rhesus macaques infected with either SIV or SHIV. Therefore, in NHP challenge models of repeated SIV/SHIV exposure, it is not practical to apply seroconversion as readout; instead, the primary readout is viremia. The highly reproducible viral outcome of the repeated oral SIV_mac251_ challenge model validates the use of viremia as readout for vaccine efficacy. However, the results of the various SHIV exposure models described here emphasize the need to define stringent criteria of infection when outcome measures other than viremia are utilized to assess vaccine efficacy.

In the current study, based on our findings in the repeated oral SIV_mac251_ exposure model, we had defined infection as two consecutive viral RNA positive time points. Applying this criterion, 1 of 7 SHIV-1157ipd3N4-exposed animals that showed a single post-challenge viral blip was considered uninfected. The remaining 6 SHIV-1157ipd3N4-exposed infants were considered infected, although one of these animals only had transient viremia. As discussed above, in human vaccine trials, weekly measurements of viremia in real time are not feasible, and viral blips as observed in 2 of 7 SHIV-1157ipd3N4-exposed infant macaques, would likely be missed. The fact that these two animals also exhibited delayed or no seroconversion, further confirmed this conclusion. In contrast, SHIV-1157ipd3N4-exposed animals with high peak viremia (>10^6^ copies/ ml) in the acute phase, seroconverted rapidly. This result is remarkably similar to the outcome of the clade C SHIV-2873Nip intrarectal challenge in adult macaques (55). The former study established that viremia below 10^4^ copies of viral RNA/ml was insufficient to result in seroconversion. Therefore, to achieve “persistent systemic infection” (PSI) in the acute phase, viral challenges should be continued until the threshold of 10^4^ copies of viral RNA/ml is reached, because a certain duration and magnitude of viremia is necessary for sufficient antigen stimulation to trigger a humoral response (55). The dose of 20-40 TCID_50_/ml of SHIV-1157ipd3N4 applied in the current study was likely a contributing factor to the transient viremia observed in two of the infant macaques. Indirectly, this is further supported by the virological outcome after repeated weekly exposure to much higher doses (>10^4^ TCID_50_/ml) of SHIV-1157(QNE)Y173H or SHIV CH505 375H.dCT. All SHIV-1157(QNE)Y173H or SHIV CH505 375H.dCT – exposed animals developed high peak viremia that fell within a narrow range among animals infected with the same virus.

Chronic SHIV viremia varied widely in all SHIV-infected groups. Among the persistently infected SHIV-1157ipd3N4, SHIV-1157(QNE)Y173H, or SHIV CH505 375H.dCT infant macaques, we could distinguish between two patterns. Some animals reached an apparent viral setpoint, while viremia continued to fluctuate in other animals. This variability in chronic viremia is reminiscent of earlier SHIVs, such as the clade B Env SHIV SF162P3 or SHIV-BaLP4 (56, 57). Our results further suggest that the modifications introduced to SHIV-1157ipd3N4, SHIV-1157(QNE)Y173H, or SHIV CH505 375H.dCT by *in vivo* passage and/or specific genetic manipulation to increase the pathogenicity in rhesus macaques may have increased peak viremia, but were less efficient in establishing persistent high-level chronic phase viremia. The lack of persistent high viremia, as is generally observed in ART-naïve HIV-infected human infants, and is recapitulated in the pediatric SIV_mac251_ rhesus macaque infection model, but not with the present SHIVs represents a limitation of these SHIV viruses. However, the variability in chronic SHIV-1157ipd3N4, SHIV-1157(QNE)Y173H, or SHIV CH505 375H.dCT viremia is also observed in adult macaques (39, 49, 58), indicating that not necessarily age, but other, additional viral and/or host-derived factors constrained the replication of the two tested SHIVs in rhesus macaques. Attenuated virus replication was also observed after intrarectal challenge of adult macaques with SHIV BG505 (59).

We want to reiterate that prevention of infection is the primary goal of an effective HIV vaccine. However, a vaccine that cannot prevent infection could potentially be of clinical importance if disease can be ameliorated and/or transmission of HIV can be significantly reduced. In fact, the analysis of the most recent human HIV vaccine trials, including the RV144 trial, added viral load as co-primary readout of vaccine efficacy (60). Therefore, we assessed secondary outcome measures of vaccine efficacy after oral SIV or SHIV exposure. Extrapolating from those data, applying a group size of 10 animals in the control and vaccine arms, and assuming at most 10 exposures per animal, each of the infection models would provide >80% power to observe an 80% reduction in the per exposure risk of infection (Table 1). Applying the stringent criterion of systemic persistent infection being defined as plasma viremia above10^4^ viral RNA copies/ ml during the acute phase, the SHIV-1157ipd3N4, SHIV-1157(QNE)Y173H, or the SHIV CH505 375H.dCT exposure model provide 99%, 98%, or 83% power, respectively, to detect 1xlog_10_ difference in peak viremia between the control and vaccine group (Table 1). The inclusion of SHIV-1157ipd3N4-infected animals with transient viremia would reduce the power to only 52% (Table 1), reiterating the importance of concisely and consistently defining infection criteria in preclinical nonhuman primate studies. Long-term virological outcome was not assessed in the current study, but the observed attenuated nature of SHIV replication suggests that viral setpoint might not be a feasible readout of vaccine efficacy in SHIV challenge models. We did instead calculate the in area-under-the curve (AUC) viremia between weeks 0 (defined as the week prior to the first viral PCR positive time point in plasma) to 8 PI, and determined that, using again group sizes of 10 animals, the SHIV-1157ipd3N4 and the SHIV CH505 375H.dCT infection models would provide 99% power to detect a 1 log_10_ difference between the control and vaccine group in AUC viremia, with somewhat lower power obtainable in the SHIV-1157(QNE)Y173H model (Table 1). In this context, it should be stressed that SIV_mac251_ is an uncloned isolate, whereas the three tested SHIVs are all clone-derived (using the SIV_mac239_ clone as background) and differ substantially in their design. Thus, caution is warranted for a direct head-to-head comparison of these viruses.

**Table 1:**
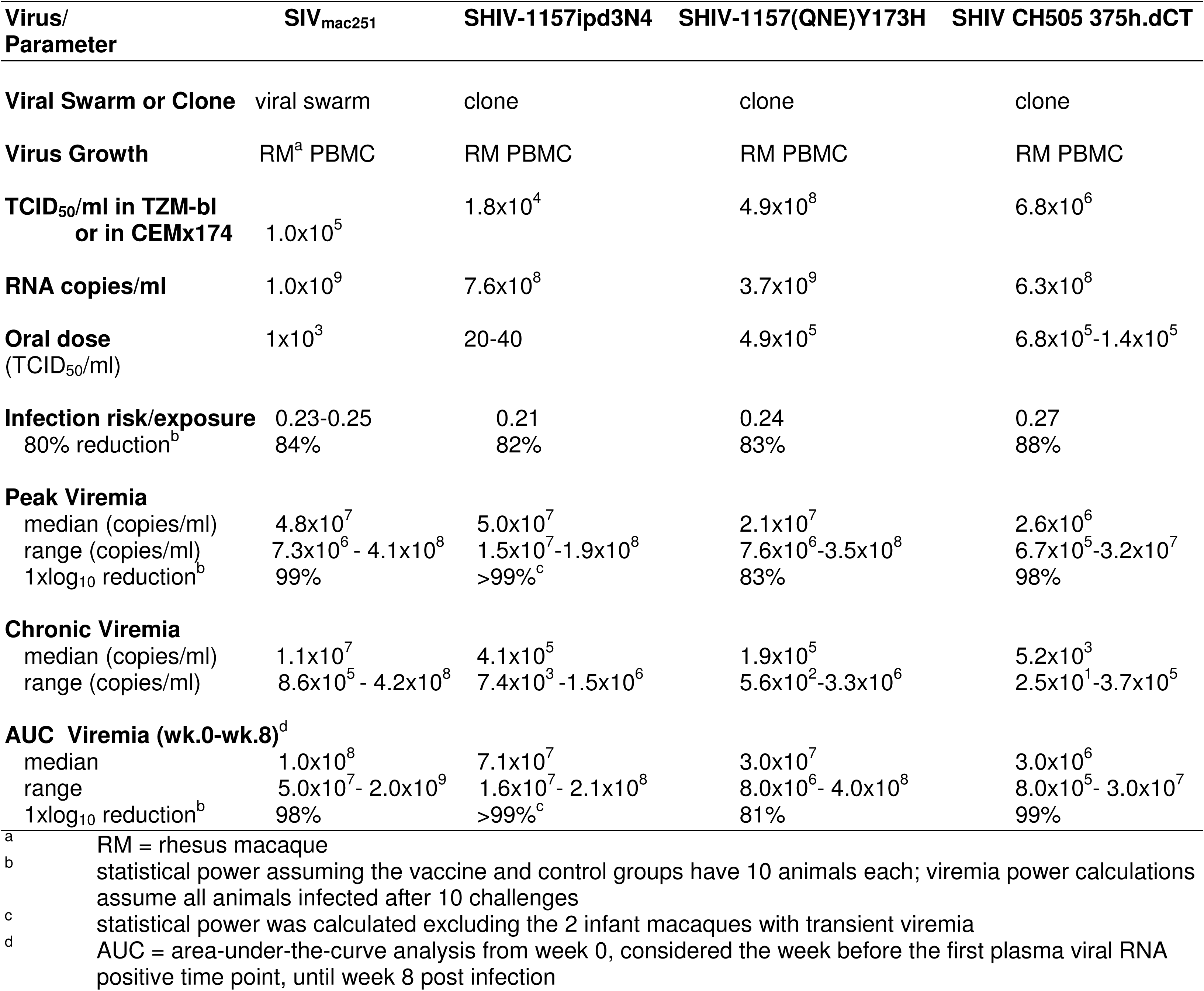
Virus Characteristics and Statistical Power Considerations.

The choice of a challenge virus and model depends on many different factors. The SIV_mac251_ infection model would be a poor choice to evaluate the role of HIV Env-directed neutralizing antibodies in vaccine-mediated protection because SIV_mac251_ is highly neutralization resistant and the SIV Env is distinct from HIV Env. In contrast, both SHIV-1157ipd3N4 and SHIV CH505 375H.dCT are considered Tier 2 neutralization sensitive viruses (40, 41). In vaccine studies testing the importance of Env-specific antibodies with Fc-mediated effector function, it might prove useful to start with challenge viruses that share a high degree of homology in specific Env epitopes, including determinants in the V1V2 and V3 regions that have been associated with protection in the RV144 HIV vaccine trial in human adults (36), before testing protection against a challenge virus heterologous to epitopes of the immunogen. Indeed, the SHIV-1157(QNE)Y173H was specifically designed to better match V2 regions of immunogens and improve recognition by neutralizing antibodies in a vaccine study (39).

Despite some of the above discussed limitations of the tested SHIV models, the pediatric oral SHIV-1157ipd3N4, SHIV-1157(QNE)Y173H, or SHIV CH505 375H.dCT rhesus macaque infection models represent a major advancement compared to earlier developed SHIVs that were not derived from clinically relevant HIV viruses. These novel clade C HIV Env SHIV challenge models will likely serve as an important resource and foundation to optimize infection models for pediatric HIV pathogenesis, vaccine, or cure studies. Still, there is need for continued development and optimization of novel SHIVs that would enable us to more readily translate findings from NHP models, including impacts on chronic phase viral replication, more reliably to human clinical studies.

## MATERIALS AND METHODS

*Animals:* SIV and type D retrovirus negative newborn Indian-origin rhesus macaques (*Macaca mulatta*) were hand-reared in the nursery of the California National Primate Research Center (CNPRC; Davis, CA) in accordance with the American Association for Accreditation of Laboratory Animal Care standards. The study strictly adhered to the guidelines outlined in The *Guide for the Care and Use of Laboratory Animals* by the Committee on Care and Use of Laboratory Animals of the Institute of Laboratory Resources, National Resource Council (61) and the *International Guiding Principles for Biomedical Research Involving Animals*. The Institutional Animal Care and Use Committee at University of California at Davis reviewed and approved the protocols prior to study initiation. Animals were anesthetized with 10 mg/kg of ketamine-HCl (Parke-Davis, Morris Plains, NJ) via the intramuscular (IM) route for oral virus challenge (see below) and sample collection.

*SIV*_*mac251*_ *infection:* The animals orally infected with SIV_mac251_ represent control animals of previously published studies (33–35). The SIVmac251 stock was propagated in rhesus PBMC. At the time of the first oral SIV_mac251_ exposure, infant macaques ranged in age from 9 weeks (n=11) (33, 34) to 12 weeks (n=6) (35). Virus was slowly applied once weekly with a needle-less syringe to the buccal mucosa in a total volume of 1 ml for a maximum of 10 exposures. Virus challenges were stopped once infection was confirmed by two consecutive plasma samples with RT-PCR-positive results (see below).

*Infection with SHIV-1157ipd3N4.* An aliquot of SHIV-1157ipd3N4 (stock 11/08/2009 with 1.9×10^4^ TCID_50_ in TZM-bl) was kindly provided by Dr. Ruth Ruprecht (Texas Biomedical Research Institute, San Antonio, TX). We prepared a virus stock by growing SHIV1157ipd3N4 in adult rhesus PBMC as described (41, 46-48), with virus growth media consisting of RPMI 1640 supplemented with 10% heat-inactivated FBS and contained *Staphylococcus aureus enterotoxin* A (SEA; Toxin Technologies, Inc; Sarasota, FL) at 0.05 μg/ml, recombinant human TNF-α (ProSpec-Tany TechnoGene Ltd., Ness Ziona, Israel) at 10 ng/ ml, and recombinant human IL-2 (ProSpec-Tany TechnoGene Ltd) at 50-100 U/ ml. An aliquot of the virus-containing culture supernatant was grown in TZM-bl cells and determined to have a titer of 18,000 TCID_50_/ml. RT-PCR analysis (see below) of multiple serial dilutions of the virus stock yielded an average value of 7.6×10^8^ RNA copies/ ml.

To determine the infectious dose in infant macaques, we used a total of 8 animals at 12 weeks of age. The infant macaques that were challenged orally with 1 ml 20 TCID_50_ of SHIV-1157ipd3N4 received their first virus exposure at 6 weeks of age and continued to be orally exposed once weekly until infection was confirmed by plasma viremia as described above for SIV_mac251_.

The related virus SHIV-1157(QNE)Y173H was kindly provided by Dr. Sampa Santra (Harvard University). In TZM-bl cells, the virus had a titer of 4.9×10^8^ TCID_50_/ml that corresponded to 3.7×10^9^ viral RNA copies/ ml (39) (Dr. Santra, personal communication). The weekly oral challenge with SHIV-1157(QNE)Y173H in infant macaques (15 weeks of age) was performed with virus diluted 1:1000 or 4.9×10^5^ TCID_50_/ml in 1 ml RPMI1640.

*Infection with SHIV CH505:* The design of SHIV CH505 has been described previously (40); the amount of virus needed to conduct all the experiments was kindly provided by Dr. George Shaw (University of Pennsylvania, Philadelphia, PA). The SHIV CH505 375H.dCT stock (12/22/2015) had the following characteristics: 178 ng/mL of p27, infectious titer of 5-7 x10^6^ IU/ml in TZM-bl cells, and a titer of 186×10^5^ IU/ ml in primary rhesus CD4^+^ T cells. All infants infected with SHIV CH505 were between 4 to 8 weeks of age at the time of the first oral exposure. Virus was administered orally in 1 ml once weekly.

*Measurement of plasma viral RNA by RT-PCR:* Plasma samples were analyzed weekly after the initiation of oral exposures using reverse transcription-PCR (RT-PCR) for SIVgag RNA as described {Li, 2016 #2355}. Note that RNA was extracted manually if plasma volumes were limited. Data are reported as the number of SIV RNA copy equivalents per ml of plasma, with a limit of detection of 15 copies/ ml. Animals with two consecutive viral RNA positive time points were considered infected and oral challenges were stopped.

*Detection of virally-infected cells in lymph node sections:* Sequential sections (5 µm) of formalin-fixed, paraffin-embedded lymph nodes were cut and stained for CD3 and CD20 as previously described (22, 35). SHIV RNA was visualized with the 1-Plex ViewRNA^™^ ISH Tissue Assay Kit using SIV_mac239_ or Beta actin (positive control) probe sets that were dectected via the ViewRNA^™^ Chromogenic Signal Amplification Kit (all from ThermoFisher, Waltham, MA). These two sequential slides were indivudually imaged with a Zeiss AxioObserver microscope and AxioCam MRm camera. Composite overlays of CD3/CD20 stained slides with ISH slides were prepared using Zen Lite v2.3 software (Zeiss).

*Plasma IgG antibody responses* were measured as previously described using gp120 of SHIV-1157 or SHIV CH505 375H.dCT, respectively (29, 50).

*CD4:CD8 T cell ratios* were determined based on Complete Blood Counts and CD4^+^ and CD8^+^ T cell percentages measured by flow cytometry (21, 28, 33, 34).

*Statistical Analysis:* Risk of infection per exposure was estimated by the number of infected macaques divided by the total number of challenges, which corresponds to the maximum likelihood estimator under certain assumptions (63) (64). Statistical power was estimated by simulation. The area-under-the curve (AUC) analysis of viremia was performed using GraphPad Prism (GraphPad, La Jolla, CA) and viremia AUC data between two groups were compared by Mann-Whitney test.

## Acknowledgements

The work was supported by National Institutes of Health grants 1R56 DE026321 (KDP), P01 AI117915 (SRP, KDP), UM1 AI100645 (GS), T32 5108303 (ADC), the Office of Research Infrastructure Programs/OD P51OD011107 (to CNPRC), and the Center for AIDS Research award P30AI050410 (to UNC). This work was supported in part with federal funds from the National Cancer Institute, National Institutes of Health, under contract no. HSN261200800001E. The content of this publication does not necessarily reflect the views or policies of the Department of Health and Human Services, nor does mention of any trade names, commercial products, or organizations imply endorsement by the U.S. government.

The content is solely the responsibility of the authors and does not necessarily represent the official views of the National Institutes of Health.

We would like to thanks Dr. Santra Sampa, Harvard Medical School, Boston, MA for providing us with SHIV1157(QNE)Y173H.

The authors thank Randy Fast, Kelli Oswald, and Rebecca Shoemaker in the Quantitative Molecular Diagnostics Core of the AIDS and Cancer Virus Program of the Frederick National Laboratory for expert assistance with viral load measurements. We would also like to thank Jennifer Watanabe, Jodi Usachenko, and the staff of the CNPRC Colony Research Services for their support in these studies.

## Author Contributions

KKAVR, SP, and KDP designed the experiments; RT, NC, MD, RG, ANN, and ADC II performed the experiments and analyzed the data; MH performed the statistical analysis; JL oversaw the viral load analysis; RMR and GS provided virus and consulted on the infection model; KKAVR, JL, SP, RMR, and KDP wrote the manuscript.

